# Spatially guided distractor suppression during visual search

**DOI:** 10.1101/2020.07.25.220947

**Authors:** Tobias Feldmann-Wüstefeld, Marina Weinberger, Edward Awh

## Abstract

Past work has demonstrated that active suppression of salient distractors is a critical part of visual selection. Evidence for goal-driven suppression includes below-baseline visual encoding at the position of salient distractors (Gaspelin and Luck, 2015) and neural signals such as the Pd that track the position and number of distractors in the visual field (Feldmann-Wustefeld and Vogel, 2019). One basic question regarding distractor suppression is whether it is inherently spatial or nonspatial in character. Indeed, past work has shown that distractors evoke both spatial (Theeuwes, 1992) and nonspatial forms of interference (Folk and Remington, 1998), motivating a direct examination of whether space is integral to goal-driven distractor suppression. Here, we provide clear evidence for a spatial gradient of suppression surrounding salient singleton distractors. Replicating past work, both reaction time and neural indices of target selection improved monotonically as the distance between target and distractor increased. Importantly, these target selection effects were paralleled by a monotonic decline in the amplitude of the Pd, an electrophysiological index of distractor suppression. Moreover, multivariate analyses revealed spatially selective activity in the theta band that tracked the position of the target and – critically – revealed suppressed activity at spatial channels centered on distractor positions. Thus, goal-driven selection of relevant over irrelevant information benefits from a spatial gradient of suppression surrounding salient distractors.

## Introduction

Visual attention allows us to cope with the vast amount of incoming information by selecting the most relevant subset of stimuli in our environment. Models of attentional control acknowledge multiple distinct influences on visual selection, including physical salience (Theeuwes, 2010), current goals (Folk & Remington, 2006), and learning experience (Awh et al., 2012; Hutchison and Turk-Browne, 2012). Goal-driven selection of targets amongst distractors has been shown to invoke a combination of target enhancement and distractor suppression. Relevant features can be enhanced to facilitate target processing (Desimone & Duncan, 1995; Wolfe, 2007) while irrelevant features can be suppressed below baseline to reduce distraction (Gaspelin et al., 2015). Neurally, task-relevant items elicit an N2pc component in the event-related potential of the EEG signal, tracking the timing and probability of visual selection (Eimer, 1996; Luck & Hillyard, 1994). In addition, salient distractors elicit a Pd component, indicative of attentional suppression of irrelevant and potentially distracting stimuli (Feldmann-Wüstefeld et al., 2020; Gaspelin & Luck, 2018; Hickey et al., 2009). Thus, recent models of visual attention implement enhancement and suppression as independent processes that resolve the competition of simultaneously presented stimuli for attentional resources (Liesefeld & Müller, 2019; Wyble et al., 2018).

A fundamental question about distractor suppression is whether it is spatial in character. Distractors can capture spatial attention. For example in visual search tasks, salient distractors elicit an N2pc, indicative of shifts of attention towards their position (Feldmann-Wüstefeld et al., 2013; Sawaki & Luck, 2013). Further, RT costs for distractors monotonically increase (Mounts, 2000) and N2pc amplitudes decrease (Gaspar & Mcdonald, 2014) when target and distractor are closer together. Thus, it is already clear that the spatial arrangement of targets and distractors has a strong influence on distractor interference. But this still leaves open the question of whether distractor suppression is minimized via suppression of visual responses in specific locations. Moreover, several studies have provided evidence for *nonspatial* filtering costs, by showing that only distractors congruent with a target template result in involuntary spatial shifts of attention whereas incongruent distractors induce RT costs irrespective of distractor location (Folk & Remington, 1998; see also Becker, 2007; Folk & Remington, 2006). Further, the presence of a salient distractor can induce longer RTs and a delayed target-N2pc without eliciting an N2pc itself (Wykowska & Schubö, 2011). Thus, there is clear evidence for nonspatial forms of distractor interference that could also be managed via distractor suppression.

The robust evidence for both spatial and nonspatial forms of interference motivates a direct examination of whether distractor suppression is spatial in character. While the contralateral nature of the Pd shows some spatial specificity, lateralized activity alone does not necessarily reflect a precise spatial representation. Thus, even with lateralized neural activity such as the Pd it is possible that suppression is not based on distractor position per se. For example, suppression might inhibit distractor processing at later stages where position is no longer critical (e.g., decision or response selection stages of processing). The present work provides evidence for spatially-directed suppression of distractors by manipulating the distance between target and distractor stimuli, and observing the consequences for behavioral and neural indices of selection and suppression. The amplitude of the Pd component covaried with the distance between target and distractor, with monotonic increases in Pd amplitude as target and distractor were presented closer together. In addition, the Pd was eliminated when target and distractor shared a location, and spatial suppression would have also negatively impacted target processing. Finally, IEM analysis of EEG oscillations (Foster et al, 2017) in the theta band tracked the target position and additionally showed a reliable depression of spatial channel activity at the distractor position, providing converging evidence for a spatial gradient of suppression surrounding salient distractors.

## Methods

### Overview

First, in two behavioral experiments we revised the additional singleton paradigm to statistically unyoke target and distractor location, i.e. to present targets and distractors in the same location in 1/d of the distractor-present trials where d = display size. This was necessary to facilitate spatial modeling of target and distractor positions, and to avoid inducing strategies that affect attentional capture (Becker, 2007; Yantis & Egeth, 1999). Furthermore, it allowed us to examine neural indices of selection and suppression when the target and distractor shared the same location, a condition that is largely absent in prior research. In singleton-present trials of Experiment 1, the target was red in 1/d of the trials while one of the nontargets was red in the remaining d-1/d of the trials. In singleton-present trials of in Experiment 2, an additional red ring was presented around the target in 1/d of the trials and around one of the nontargets in d-1/d of the trials (i.e., one of the items was a triple-compound stimulus). We then used the triple-compound variant in two EEG experiments. In Experiment 3, the distractor consistently had the same color throughout the experiment whereas in Experiment 4, it had a different color when it was presented in the target location, which allowed observers to rely on color as a defining property of the target. As behavioral results were virtually identical in Experiment 3 and 4, we collapsed across these experiments to achieve a higher statistical power for EEG analyses.

### Participants

In Experiment 1, twenty volunteers aged 19-31 years (*M* = 21.5, *SD* = 3.2), thirteen female, all right-handed, participated. In Experiment 2, twenty-one volunteers aged 18-31 years (*M* = 23.5, *SD* = 4.5), eleven female, one left-handed participated. In Experiment 3, participants were aged 18-31 years (*M* = 22.8, *SD* = 4.0), nine participants were female, and five were left-handed. In Experiment 4, participants were aged 18-31 years (*M* = 22.8, *SD* = 4.0), nine participants were female, and five were left-handed. In Experiment 4, participants were aged 18-33 years (*M* = 22.2, *SD* = 3.7), sixteen participants were female, and one was left-handed. Participants were naive to the paradigm and objectives of the study and were paid ~$10 (Experiment 1 and 2) or ~$15 (Experiment 3 and 4) per hour. All experiments were conducted with the written understanding and consent of each participant. The University of Chicago Institutional Review Board approved experimental procedures.

### Apparatus

Participants were seated in a dimly lit, electrically shielded and sound attenuated chamber. Participants responded with button presses on a standard keyboard that was placed in front of them. Stimuli were presented on an LCD computer screen (BenQ XL2430T; 120 Hz refresh rate; 61 cm screen size in diameter; 1920 × 1080 pixels) placed at 75 cm distance from participants. An IBM-compatible computer (Dell Optiplex 9020) controlled stimulus presentation and response collection using MATLAB routines. In Experiment 3 and 4, participants were seated with a chin-rest to allow for tracking gaze position.

### Stimuli

All stimuli were presented on a gray background (RGB: 128-128-128). Search displays always showed six items presented equidistantly on an imaginary circle (3° radius) around a black central fixation cross (0.5° diameter, RGB: 0-0-0), see Fig. 1. Items positions were fixed with two items on the vertical midline and two items in each hemifield. One of the items was a diamond (2.1° diameter, 2 pixels line width), serving as a target stimulus. The other five items were nontarget circles (1.6° diameter, same area as target, 2 pixels line width, RGB: 64-64-64). In distractor-absent trials, all items were dark grey (RGB: 64-64-64) and the target appeared equally often at one of the six positions. In distractor-present trials of Experiment 1, one of the six items was red (RGB: 255-0-0). The singleton was equally often presented in each location, i.e., the target was a red diamond in 1/6 of the distractor-present trials. In distractor-present trials of Experiment 2 and 3, an additional empty red circle (2.4° diameter, 2 pixels line width, RGB: 255-0-0) appeared equally often around any one of the six items, i.e., around the target in 1/6 of the distractor-present trials. In Experiment 4, the distractor had distinct colors for target versus nontarget positions (see Fig. 1C). Distractor were red/green for nontarget locations and green/red for target locations (balanced across participants). The 36 combinations of target and distractor locations appeared equally often in distractor-present trials. Within each of the six items was a dark gray line that was either vertical or horizontal (0.75° length, RGB: 64-64-64). Horizontal and vertical lines were used equally often for the target. In addition, in distractor-present trials in which the distractor was presented at a location different from the target location, the four combinations of horizontal and vertical lines were used equally often for the target and the distractor. The orientation of the lines within the remaining nontargets was randomly assigned with as many horizontal as vertical lines (or as close as possible).

**Figure 1.**
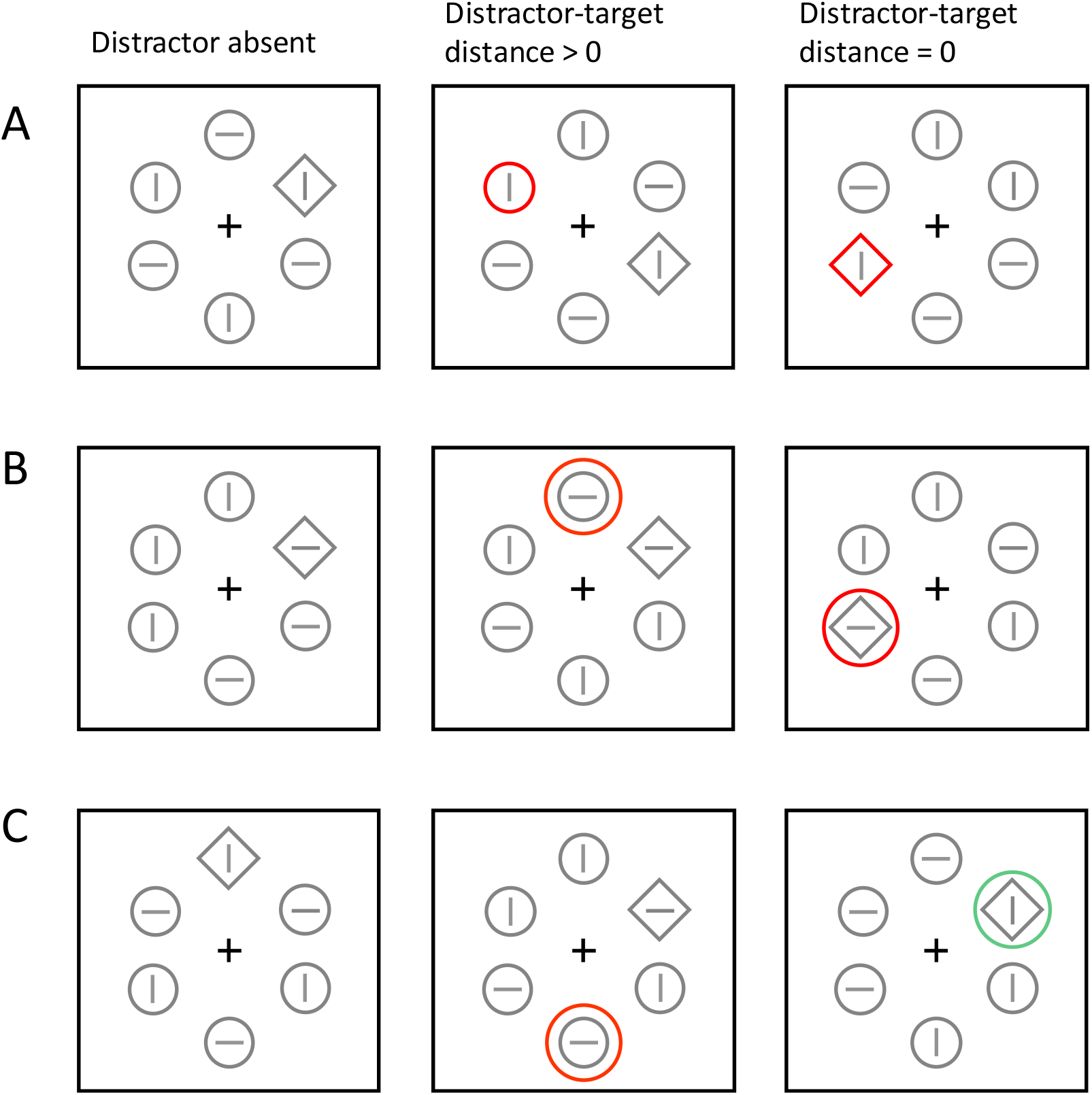
Visual search displays used for Experiment 1 **(A)**, Experiment 2 & 3 **(B)** and Experiment 4 **(C)**. Participants had to find the diamond-shaped target and report the orientation of its embedded line with a key press. In half the trials, all items were grey (distractor absent trials). In the other half of the trials, a color singleton (red/green ring), serving as a distractor, was presented equally likely in one of the six positions. This means the singleton was in a distractor position in 5/6 of singleton-present trials (distance > 0) and in a target position in 1/6 of singleton-present trials (distance = 0). In Experiment 4, different color were used for distance > 0 and distance = 0 trials.

**Figure 2.**
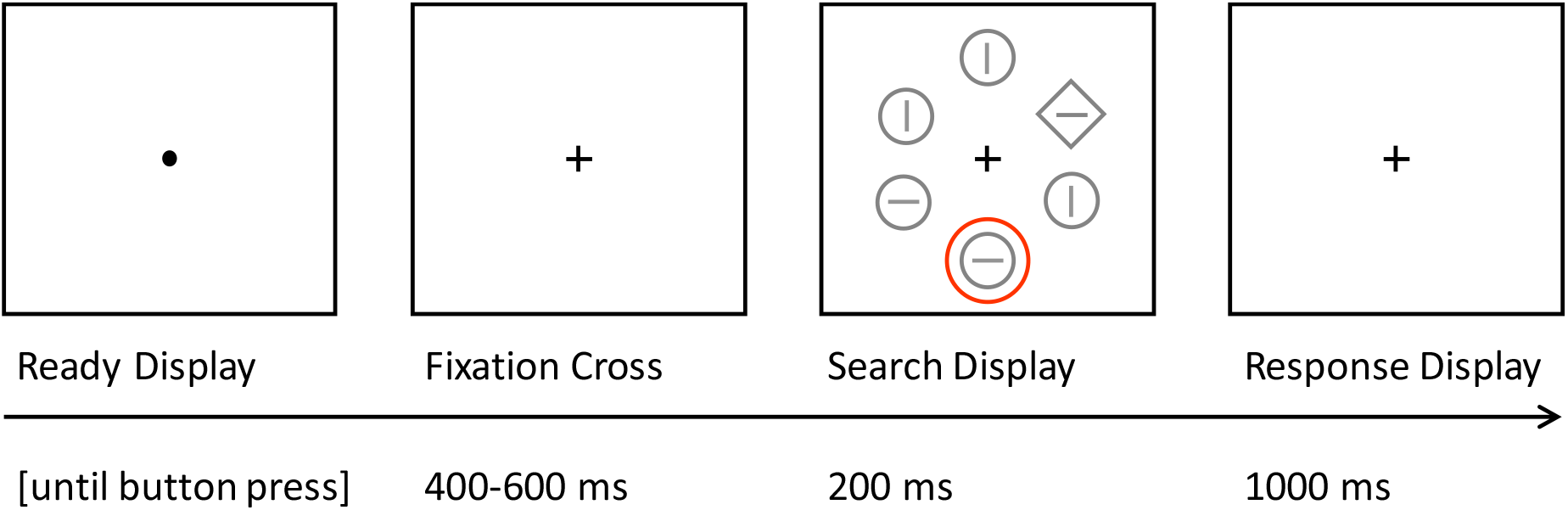
Trial procedure. Each trial started with a Ready Display. Participants had to fixate the central dot and press a key when ready. Upon key press a fixation cross followed for 400-600 ms before the search display was presented for 200 ms. The subsequent Response display only showed a fixation cross. Participants had to respond during the Search Display or Response Display presentation (i.e., within 1200 ms). The Response display was followed by an inter-trial interval showing a blank screen for 1000 ms (Experiment 1) or 800-1200 ms (Experiment 2&3). The Ready Display was not used in Experiment 1 & 2.

### Procedure

Each trial started with a display showing the fixation cross and no other objects for a duration of 500 ms (Experiments 1 and 2) or a randomly varying duration between 400 and 600 ms (Experiments 3 and 4). Subsequently, the search display was shown for 200 msec in addition to the fixation cross. Finally, a response display with the fixation cross only was shown until participants responded or for a maximum of 1000 msec. Participants were instructed to report the orientation (horizontal or vertical) of the line within the diamond-shaped target by pressing a key corresponding to each orientation (key assignments were counterbalanced across participants). Their responses were limited to a time window of 1200 msec (search display + response display), and responses made after this period were not counted. If participants did not respond within this time window, a message reading “Too slow” was shown for 500 msec. After this message, or after participants responded, a blank screen was shown for a duration of 1000 msec (Experiments 1 and 2) or a randomly varying duration between 800 and 1200 ms (Experiments 3 and 4). To optimally prepare participants to avoid eye movements which would abort a trial (see below), a preparation display was shown prior to the start of a trial in Experiments 3 and 4. The preparation display showed a single black fixation dot in the center of the screen (0.25° diameter, RGB: 0-0-0). Participants were told to fixate the dot and not move their eyes or blink until the end of a trial. After subjects pressed the spacebar, the trial started with showing the fixation cross.

Participants were given the opportunity to practice the task before the experiment began. Once participants reached 75% accuracy and felt comfortable doing the task, they began the experiment.

Experiments 1 and 2 consisted of 24 blocks of 36 trials (864 trials in total). Half the trials (432) were singleton-absent, the other half (432) were singleton-present. Experiments 3 and 4 consisted of 28 blocks of 72 trials (2016 trials in total). Half the trials (1008) were distractor-absent, the other half (1008) were distractor-present. All trials were presented in random order across an entire experiment. After each block, participants were given feedback about their performance (percent correct, number of slow trials, and if a new “high score” was achieved for accuracy during that block). Following feedback, there was a mandatory ten second break after which participants could choose to either immediately continue to the next block or to extend their break.

### Eye tracking (Experiments 3 and 4 only)

In Experiments 3 and 4, gaze position was tracked at a sampling rate of 1000 Hz for both eyes with an EyeLink 1000+ eye tracker (SR Research Ltd.). A direct gaze violation feedback procedure was applied from halfway through the fixation display until participants responsed or the response time window ended. The trial was aborted if the participants’ gaze was not within 1.5° of the center of the fixation cross during this time or if they blinked. A message (either “Eye movement” or “Blink”) was presented on the screen, and then the next trial began with the new preparation display. Each aborted trial was added back into the sequence of upcoming trials, extending the experiment by one trial, and the trial sequence was re-shuffled so as to make the aborted trial’s reappearance unpredictable. The mean number of total trials (including repeated trials) was 2140 (SD = 126) in Experiment 3 and 2224 (SD = 171) in Experiment 4.

### EEG Recording (Experiments 3 and 4 only)

EEG was recorded using Ag-AgCl active electrodes (Brain-Products actiCAP) from 32 scalp sites (according to the International 10/20 System: FP 1/2, F7/8, F3/4, Fz, FC5/6, FC1/2, C3/4, Cz, TP9/10, CP5/6, CP1/2, P7/8, P3/4, PO7/8, PO3/4, Pz, O1/2, Oz). FPz served as the ground electrode and all electrodes were referenced online to TP10 and re-referenced off-line to the average of all electrodes. Incoming data were filtered [low cut-off=.01 Hz, high cut-off = 250 Hz, slope from low-to high-cutoff = 12 dB/octave] and recorded with a 1000 Hz sampling rate. Impedances were kept below 10 kΩ. Horizontal and vertical electrooculograms (EOGs) were recorded with passive electrodes bipolarly of approximately 1 cm from the outer canthi of the eyes and from 1 cm above and 2 cm below the observers’ right eye, respectively.

### Design & data analysis

There were two main conditions: distractor-absent and distractor-present trials. Since trials with coinciding target and distractor constitute a special case, three conditions were considered: distractor-absent, distractor present, distractor-target distance = 0 (from here on “Dist=0”), distractor-target distance > 0 (from here on “Dist>0”). Accordingly, mean response times (RTs) and error rates were forwarded to a one-way ANOVA for repeated measures with three factor levels. In addition, RTs in distractor-present trials were analyzed as a function of distance between target and distractor. As there were six positions and target and distractor could be presented at the same location, there was a total of four distances used (0, 1, 2, 3). For all RT analyses, only correct trials were used. For all experiments, Greenhouse-Geisser correction was applied where appropriate. Effect sizes are reported as partial eta squared (η^2^) for ANOVAs and Cohen’s d for t-tests.

Experiments 3 and 4 had the same conditions regarding behavioral measures as Experiments 1 and 2. With regards to event-related potentials (ERPs), the lateralization of stimuli plays a crucial role (Hickey et al., 2009; Woodman & Luck, 2003). For ERP analyses, we focused on trials in which either the target was presented laterally and the distractor was on the vertical midline (or absent) or the distractor was presented laterally and the target was on the vertical midline. These trials do not confound target and distractor activity in the lateralized ERP and allow to disentangle target and distractor processing. The ERP was analyzed separately for target-distractor distances 0, 1, or 2. This resulted in seven conditions for ERP analyses in both Experiment 3 and 4: (i) Target lateral, singleton absent; (ii) Target and singleton at same lateral location, distance 0; (iii) Target lateral, singleton midline, distance 1; (iv) Target lateral, singleton midline, distance 2; (v) Singleton lateral, target midline, distance 1; (vi) Singleton lateral, target midline, distance 2.

### EEG Data Analysis (only Experiments 3 and 4)

EEG was averaged off-line over a 700-ms epoch including a 200-ms pre-stimulus baseline with epochs time-locked to search display onset. Trials with incorrect or no responses were excluded. Trials with eye-related artifacts from −200 to 600 ms were excluded from the analysis (Experiment 2: M = 18.0%, SD = 11.3%; Experiment 3: M = 23.8%, SD = 12.8%). To identify eye-related artifacts, eye tracking data was first baselined identically to EEG data (i.e., subtraction of the mean amplitude of x and y coordinates for the time from −200 – 0 ms). Then, the Euclidian distance from the fixation cross was calculated from baselined data. We identified saccades with a step criterion of 0.5° (comparing the mean position in the first half of a 50-ms window with the mean position in the second half of a 50-ms window; window moved in 20-ms steps). We identified drifts by eye tracking data indicating a distance from the fixation > 1.2°. To minimize false alarms due to noise in the data, both eyes had to indicate an eye-related artifact for a trial to be excluded from analysis. For one participant, eye tracking data of sufficient quality was not available. For this participant, the same procedure as for eye tracking based artifact rejection was used with these criteria: 25 μV hEOG (difference left and right channel), 100 μV for vEOG (difference upper and lower channel). Participants with fewer than 1000 trials were excluded from analyses (Experiment 3: n = 1; Experiment 4: n = 2).

#### Event-related potentials (ERPs)

For all ERP-relevant conditions (see above), mean contralateral and ipsilateral ERP activity was calculated for each participant for an electrode pool of the three symmetric sites PO7/PO8, P7/8, PO3/4. For statistical analyses, the lateralized waveform (contra- minus ipsilateral activity) was used. The time windows for statistical analyses were determined in a data-driven way. For each component of interest, a 100-ms time window was determined around the peak of the component collapsed across conditions. The mean amplitude of the respective component was calculated for these time windows and forwarded to ANOVAs for repeated measures. The N2pc was compared for trials with lateral targets and no distractor, distractors at distance 1 and distractors at distance 2. The Ppc, distractor-N2pc, and Pd were compared for trials with lateral distractors and targets in the same position as the distractor, for targets at distance 1, and for targets at distance 2.

#### Inverted encoding model

While ERPs allow the measurement of visual priority on the coarse scale of left versus right hemifield, inverted encoding models can be used to reconstruct spatially selective channel tuning functions from the topographic distribution of EEG activity across electrodes (Ester et al., 2015; Fahrenfort et al., 2016; Foster et al., 2016). We used an IEM that assumes that power measured at each electrode reflects the weighted sum of six spatial channels (representing neuronal populations), each tuned for one of the six item positions. The IEM was run on evoked power (activity phase-locked to stimulus onset). To isolate frequency-specific activity, raw EEG segments were band-pass filtered for the alpha frequency range (8-12 Hz) using a two-way least-squares finite-impulse-response filter (‘‘eegfilt.m’’ from EEGLAB Toolbox) and then hilbert-transformed. The complex analytic signal was first averaged across trials, and then the complex magnitude of the averaged analytic signal was squared. In a training stage, segmented, filtered waveforms from distractor-absent trials (equaled for target location) were used to estimate the weights for a model of the target’s spatial position in a least-squares estimation. Each artifact-free trials were assigned to one if two bins (block assignment had 1000 iterations to allow for cross-validation). In the test stage, the model was inverted to transform the remaining half of trials (equaled for positions) into estimated channel responses, using the previously determined weights, for each of the two blocks. Channel responses were averaged across blocks and block assignment iterations. Testing was done twice using the same trials; once using the target position labels, once using the distractor position labels. This means that a common set of training data was used to estimate the channel responses for targets and singletons separately, but in the same trials. This procedure precludes the possibility that any observed effects of attention deployment are due to differences that arose during training or because of using distinct “basis sets” for the two conditions. The six channel response functions were shifted to a common center and averaged to obtain the CTF. The slope of the CTFs (estimated by linear regression computed for each time point) was used as a metric to compare attention deployment towards target and distractor. The slope was compared between channels tested on target versus distractor position with t-test for dependent measures for each time point between 0 and 500 ms. Only clusters of significance in which 50 consecutive data points (i.e., 50 ms) have to show a p < .05 were considered reliable.

## Results

### Experiment 1

Distractor-absent trials were fastest (555 ms), followed by Dist=0 trials (566 ms) and Dist>0 trials (582 ms), see upper panel in Fig. 3A. This was confirmed by a reliable main effect of Trial Type, F(2,38) = 23.7, p < .001, η^2^ = .56. All direct comparisons were significant (all p ≤ .012). Trial type also affected accuracy (M_absent_ = 90.7%, M_dist=0_ = 90.8%, M_dist>0_ = 88.6%), F(2,38) = 3.3, p = .047, η^2^ = .15. Direct comparisons show reliable differences between distractor-absent and Dist>0 trials (p = .033), but not between any other condition (both p ≥ .053).

**Figure 3.**
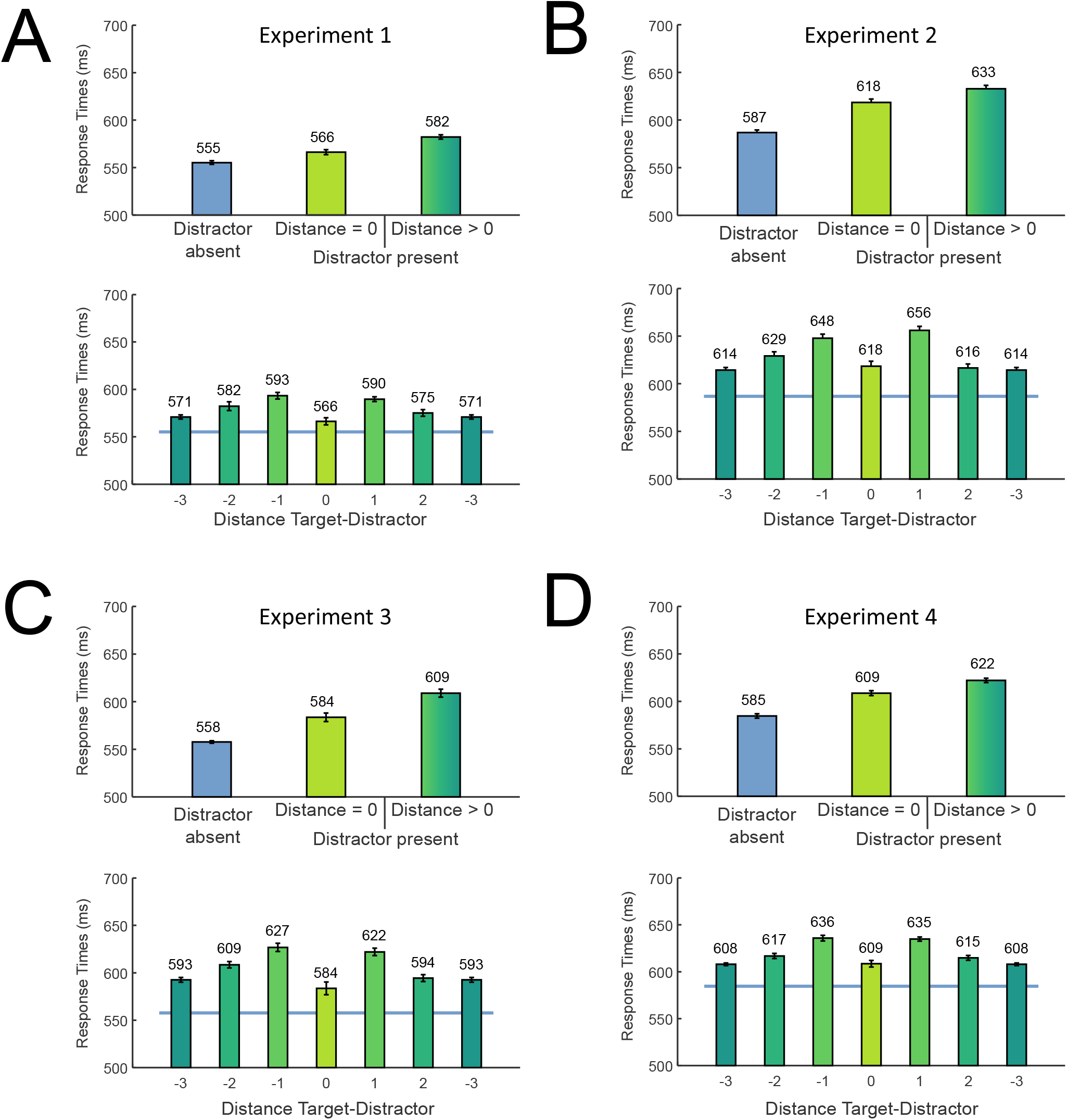
Behavioral results Experiment 1-4. The upper panels shows RT as a function of trial type (distractor absent, distractor-target distance = 0, distractor-target distance > 0). The lower panels show RT as a function of distance between target and distractor in distractor-present trials (blue line depicts RT in distractor-absent trials for comparison). Negative numbers represent counterclockwise distances, positive numbers clockwise distances. Distance 0 refers to trials in which target and distractor appeared at the same location (Experiments 2-4) or the target was the color singleton itself (Experiment 1). Error bars denote standard errors of the mean, corrected for within-subjects variability (Cousineau, 2005).

**Figure 4.**
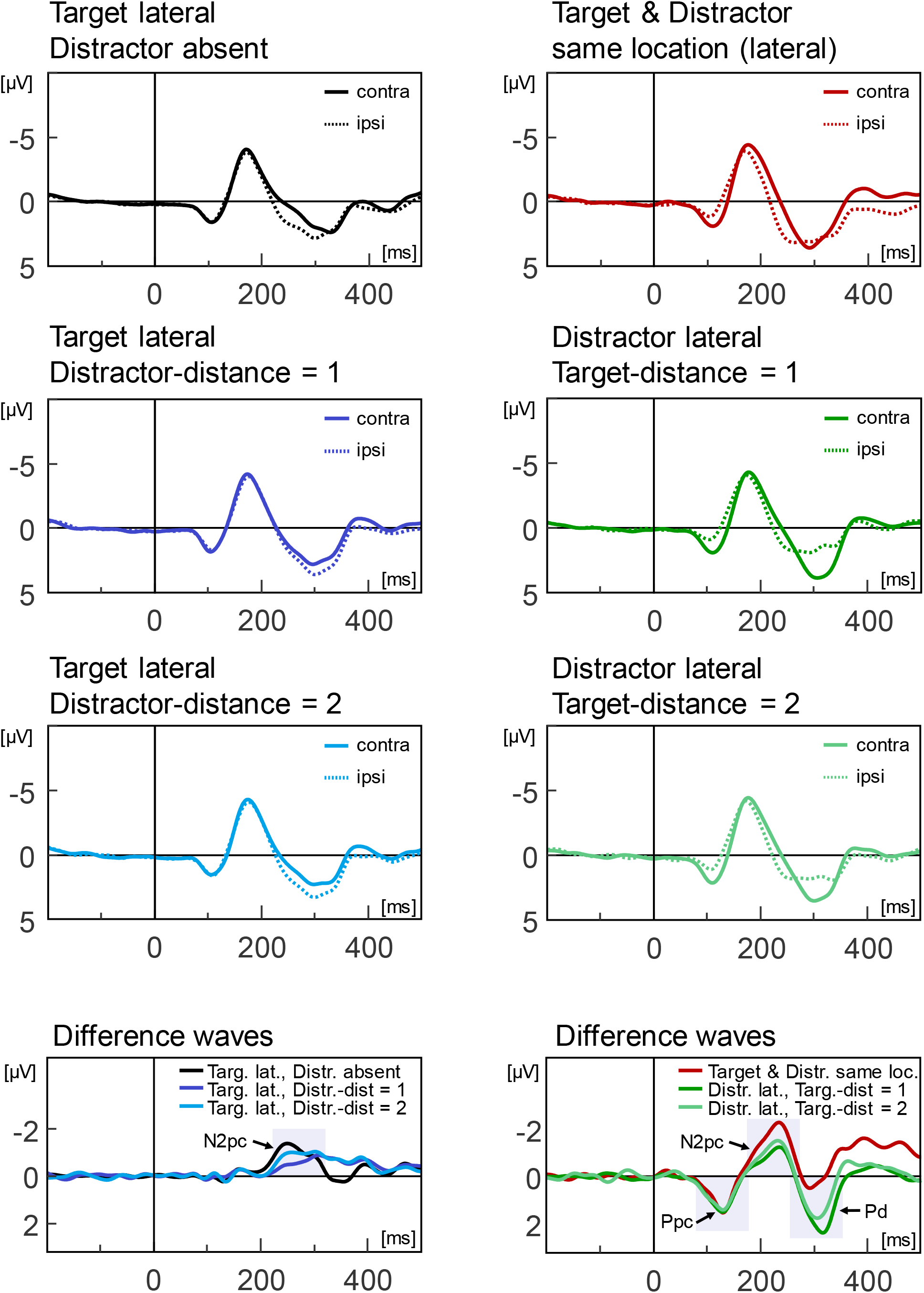
Grand average event-related potentials collapsed across Experiment 3 and 4. The upper left column shows trials in which the target was presented laterally and the distractor was presented on the vertical midline. The right panels show trials in which the distractor was presented laterally and the target was presented on the vertical midline (or target was at distractor location, red lines). The lowest row shows the difference waves (contra minus ipsilateral) for the same conditions as shown in the upper rows. Signal is pooled across PO7/8, P7/8 and PO3/4. For display purposes, signal was filtered with a 30Hz lowpass filter. All statistical analyses were conducted on unfiltered data. Grey shaded areas show the time windows usedfor statistical purposes.

In distractor-present trials, RT was fastest for dist=0 trials. For dist = 1, RT was slowest, with monotonically decreasing RTs for increasing distances (see lower panel in Fig. 3A). This was confirmed by a reliable main effect of Distance, F(3,57) = 13.7, p < .001, η^2^ = .42. All direct comparisons between distances were reliable (all p ≤ .020) except for distance 0 and 3 (p = .382). The distance between target and distractor also affected accuracy (M_0_ = 90.8 %, M_1_ = 87.9 %, M_2_ = 88.6 %, M_3_ = 90.0 %), F(3,57) = 2.9, p = .041, η^2^ = .13. Only the direct comparisons between distance 0 and 1 was reliable (p = .030), none of the other direct comparisons were reliable (both p ≥ .051).

In compatible trials (target and distractor line of same orientation) RTs were equally long (M = 581 ms) as in incompatible trials, (M = 584 ms), t(19) = 0.7, p = .236, d = 0.04. Accuracy, however, was higher in compatible (M = 89.5%) than in incompatible trials (M = 87.7%), t(19) = 1.9, p = .040, d = 0.18.

In sum, the results show best performance in distractor absent trials and a spatial gradient of interference in distractor present trials. RT rose monotonically as the distance between target and distractor declined. The compatibility effect is further evidence for spatial interference as it suggests that distractors were processed to some extent (Becker, 2007).

### Experiment 2

Similar to Experiment 1, distractor-absent trials were fastest (587 ms), followed by Dist=0 trials (618 ms) and Dist>0 trials (633 ms), see upper panel in Fig. 3B. This was confirmed by a reliable main effect of Trial Type, F(2,40) = 33.8, p < .001, η^2^ = .63. All direct comparisons were significant (all p ≤ .042). Trial type also affected accuracy (M_absent_ = 93.3%, M_dist=0_ = 91.8%, M_dist>0_ = 90.7%), F(2,40) = 4.5, p = .017, η^2^ = .18. Direct comparisons show reliable differences between distractor-absent and Dist>0 trials (p = .006), but not between any other condition (both p ≥ .073).

Similar to Experiment 1, RT was fastest for dist=0 trials and slowest for dist = 1, with monotonically decreasing RTs for increasing distances (see lower panel in Fig. 3B). This was confirmed by a reliable main effect of Distance, F(3,60) = 18.7, p < .001, η^2^ = .48. All direct comparisons between distances were reliable (all p ≤ .025) except for distance 0 and 3 (p = .553) and distance 0 and 2 (p = .511). The distance between target and distractor did not affect accuracy (M_0_ = 91.8 %, M_1_ = 89.7 %, M_2_ = 91.4 %, M_3_ = 91.3 %), F(3,60) = 1.6, p = .196, η^2^ = .08. No direct comparisons between distances was reliable (all p ≥ .097).

In compatible trials, RTs were shorter (M = 625 ms) than in incompatible trials, (M = 641 ms), t(19) = 3.8, p < .001, d = 0.20. Analogously, accuracy was higher in compatible (M = 92.3%) than in incompatible trials (M = 89.1%), t(19) = 3.0, p = .003, d = 0.53.In sum, the spatial gradient of interference as well as the compatibility effect from Experiment 1 were replicated.

### Behavioral results Experiment 3

Similar to Experiment 1 and 2, distractor-absent trials were fastest (558 ms), followed by Dist=0 trials (583 ms) and Dist>0 trials (609 ms), see upper panel in Fig. 3C. This was confirmed by a reliable main effect of Trial Type, F(2,46) = 33.7, p < .001, η^2^ = .59. All direct comparisons were significant (all p ≤ .004). Trial type also affected accuracy (M_absent_ = 92.1%, M_dist=0_ = 91.1%, M_dist>0_ = 88.9%), F(2,46) = 15.1, p < .001, η^2^ = .40. Direct comparisons show reliable differences between distractor-absent and Dist>0 trials and between distractor-absent and Dist=0 trials (both p ≤ .007), but not between Dist=0 and DIst>0 (p = .073).

Similar to Experiment 1 and 2, RT was fastest for dist=0 trials and slowest for dist = 1, with monotonically decreasing RTs for increasing distances (see lower panel in Fig. 3C). This was confirmed by a reliable main effect of Distance, F(3,69) = 13.4, p < .001, η^2^ = .37. All pairwise comparisons were significant (all p ≤ .037) except the comparison between distance 0 and 3 (p = .237). The distance between target and distractor also affected accuracy (M_0_ = 91.1 %, M_1_ = 87.2 %, M_2_ = 89.7 %, M_3_ = 90.4 %), F(3,69) = 11.8, p < .001, η^2^ = .34. Pairwise comparisons were significant (all p < .001) except for distance 0 versus 3, 0 versus 2, and 2 versus 3 (p ≥ .067).

In compatible trials (target and distractor line of same orientation) RTs were shorter (M = 596 ms) than in incompatible trials, (M = 622 ms), t(23) = 8.3, p < .001, d = 0.48. Analogously, accuracy was higher in compatible trials (M = 91.7%) than in incompatible (M = 86.1%) trials, t(23) = 5.2, p < .001, d = 0.77. In sum, the spatial gradient of interference as well as the compatibility effect from Experiment 1 and 2 were replicated.

### Behavioral results Experiment 4

Similar to Experiments 1 – 3, distractor-absent trials were fastest (585 ms), followed by Dist=0 trials (609 ms) and Dist>0 trials (622 ms), see upper panel in Fig. 3D. This was confirmed by a reliable main effect of Trial Type, F(2,44) = 41.9, p < .001, η^2^ = .66. All direct comparisons were significant (all p ≤ .005). Trial type also affected accuracy (M_absent_ = 92.7%, M_dist=0_ = 91.5%, M_dist>0_ = 90.3%), F(2,44) = 9.1, p = .001, η^2^ = .29. Direct comparisons show reliable differences between distractor-absent and Dist>0 trials and between distractor-absent and Dist=0 trials (both p ≤ .021), but not between Dist=0 and Dist>0 (p = .068).

Similar to Experiments 1 – 3, RT was fastest for dist=0 trials and slowest for dist = 1, with monotonically decreasing RTs for increasing distances (see lower panel in Fig. 3D). This was confirmed by a reliable main effect of Distance, F(3,66) = 25.7, p < .001, η^2^ = .54. All pairwise comparisons were significant (all p ≤ .002) except for the comparison distance 0 versus 3 and 0 versus 2 (both p ≥ .116). The distance between target and distractor also affected accuracy (M_0_ = 91.5 %, M_1_ = 89.1 %, M_2_ = 91.0 %, M_3_ = 91.6 %), F(3,66) = 6.1, p = .004, η^2^ = .22. Pairwise comparisons were significant (all p < .005 except for distance 0 versus 2, 0 versus 3, and 2 versus 3 (all p ≥ .285).

In compatible trials RTs were shorter (M = 615 ms) than in incompatible trials (M = 629 ms), t(22) = 5.5, p < .001, d = 0.25. Similarly, accuracy was higher (M = 92.3%) in compatible than in incompatible trials (M = 88.4%), t(22) = 6.1, p < .001, d = 0.68.

In sum, the spatial gradient of interference as well as the compatibility effect from Experiments 1 – 3 were replicated. Of particular interest for analyzing EEG data is the identical pattern of behavioral results in Experiments 3 and 4.

#### Comparison Experiment 3 and 4

The similarity of behavioral results in Experiments 3 and 4 was first confirmed with a ANOVA for repeated measures using the within-subjects factor Trial type (distractor-absent, dist=0, dist>0) and the between-subjects factor Experiment (3 versus 4), which showed neither a main effect of Experiment, F(1,45) = 2.0, p = .162, η^2^ = .04, nor an interaction of Trial type and Experiment, F(2,90) = 1.9, p = .163, η^2^ = .04. Second, an ANOVA for repeated measures was run using the within-subjects factor Distance (0, 1, 2, 3) and the between-subjects factor Experiment (3 versus 4). Neither Experiment, F(1,45) = 1.1, p = .303, η^2^ = .02 nor the interaction of Experiment and Distance, F(3,135) = 1.2, p = .293, η^2^ = .03 showed reliable effects. Thus our results indicate that using a distinct color for distance-zero trials where targets and distractor positions coincide does not yield an attention deployment different from using the same color for distance-zero and other distance > 0 trials.

In sum the behavioral results from Experiments 1-4 revealed both spatial and nonspatial filtering costs. Faster RTs with increasing distance between target and distractor indicate a spatial gradient of interference. Further evidence for spatial interference comes from the finding that distractors appearing at the same location as the target yield faster RTs than distractors appearing at different locations; when target and distractors spatially coincided, more pronounced attention deployment was observed. Moreover, compatibility effects support the notion of spatial interference; faster responses for identical target and distractor identity suggest that attention is deployed toward the distractor prior to the target in a subset of trials. Evidence for nonspatial interference comes from the comparison of distractor-absent trials and trials in which target and distractor coincide. As trials without any distractor yielded faster RTs than trials in which target and distractor appear at the same location, our results suggests that the mere presence of a distractors results in filtering costs that are not well explained by spatial capture. A pure spatial account of attentional capture (Theeuwes, 2010) would have predicted that target and distractor at the same location result in the most efficient attention deployment as two singletons coincide. Even when top-down control is facilitated by presenting the additional singleton in a distinct color when presented at the target location (Experiment 4), this effect was replicated, demonstrating robust nonspatial filtering costs.

### ERP results Experiment 3 and 4

As behavioral results were virtually identical in Experiments 3 and 4, we collapsed the EEG data from Experiments 3 and 4 to achieve a higher statistical power.

#### N2pc in target-lateral trials (223-322 ms)

The N2pc was significantly different from zero in all conditions (all p < .001). The N2pc was larger for distractors at distance 2 (M = −0.85) than at distance 1 (M = −0.56), t(46) = 2.6, p = .012, d = 0.39, suggesting more attention deployment towards the target when the distractor was further away. The N2pc was also larger in distractor-absent trials (M = −0.93) than for distractors at distance 1, t(46) = 3.2, p = .002, d = 0.51, but not larger than for distractors at distance 2, t(46) = 1.0, p = .327, d = 0.11. This suggests that a target alone was attended more strongly than a target with a nearby distractor, but not more than a target with a further away distractor. The direct comparisons of target-N2pc amplitudes between Trial Types were confirmed with a significant one-way ANOVA, F(2,92) = 7.0, p = .001, η2 = .13.

#### N2pc in distractor-lateral trials (175-274 ms)

The N2pc was significantly different from zero in all conditions (all p < .001). Distractor-N2pc amplitude was equally large for targets at distance 2 and at distance 1, t(46) = 1.7, p = .095, d = 0.16, suggesting that distractors capture attention equally strongly regardless of the distance to a target. The N2pc elicited by target and distractor in coinciding positions (M = −1.43) was larger than the distractor-N2pc for targets at distance 1 (M = − 0.69), t(46) = 6.2, p < .001, d = 0.68, and for targets at distance 2 (M = −0.85), t(46) = 5.6, p < .001, d = 0.49. In addition, the N2pc was larger in trials with coinciding target and distractor positions than in distractor absent trials, t(46) = 4.1, p < .001, d = 0.47. This indicates that attention is most strongly deployed to a location shared by target and distractor. The direct comparisons of distractor-N2pc amplitudes between Trial Types were confirmed with a significant one-way ANOVA, F(2,92) = 26.7, p < .001, η2 = .37.

#### Distractor positivity (Pd; 255-354 ms)

The Pd was significantly different from zero for distance 1 and 2 (both p < .001), except for coinciding target and distractor positions (p = .310).

The effect of distance on Pd amplitude was the opposite of the effect on the target-N2pc: Pd amplitude was larger for targets at distance 1 than at distance 2, t(46) = 3.6, p = .001, d = 0.44. At the same time, the Pd elicited by target and distractor in coinciding positions (M = −0.20) was smaller than the Pd for targets at distance 1 (M = 1.24), t(46) = 8.7, p < .001, d = 1.18, and for targets at distance 2 (M = 0.82), t(46) = 6.2, p < .001, d = 0.92. The direct comparisons of Pd amplitudes between Trial Types were confirmed with a significant one-way ANOVA, F(2,92) = 48.0, p < .001, η2 = .51.

#### Exploratory analysis: Posterior positivity contralateral (Ppc; 79-178 ms)

The Ppc was significantly different from zero in all conditions (all p < .001). Ppc amplitude did not vary as a function of distractor condition, F(2,92) = 2.2, p = .121, η2 = .05.

#### Exploratory analysis: Contralateral Delay Activity (CDA; 400-500 ms)

Visual inspection of the lateralized ERP waveforms showed that after typical analysis time windows of attentional components, a sustained negativity was found. A one-way ANOVA using each of the six conditions as a factor level indicated that CDA varied between conditions, F(5,230) = 13.7, p < .001, η2 = .23. Follow-up t-tests for dependent measures showed that the CDA elicited in trials with coinciding target and distractor positions was larger than in any other condition (all p < .001), whereas the CDA did not vary among any other conditions (all p ≥ .169), all two-tailed.

### IEM results Experiment 3 and 4

Similar to ERP analyses, we collapsed the EEG data from Experiments 3 and 4 to achieve higher statistical power. The channel tuning functions (CTFs) are shown in Fig. 5. As a measure of spatial selectivity, the slope for the channel tuning functions (the rate of change in activity level from the central target position to further away locations) was calculated for both the target location (blue lines) and the distractor location (red lines). The IEM run on alpha bandpass (8-12 Hz) filtered data shows a slope reliably different from zero from 17-500 ms (targets) and 33-289 ms (distractors). The distractor slope was reliably steeper than the target slope from 156-227 ms. The target slope was reliably steeper than the distractor slope from 328-500 ms. This shows that the channels were first selective for both target and distractor position, but more so for distractor position. Then, channels continued to be selective for the target position and not for the distractor position anymore.

**Figure 5.**
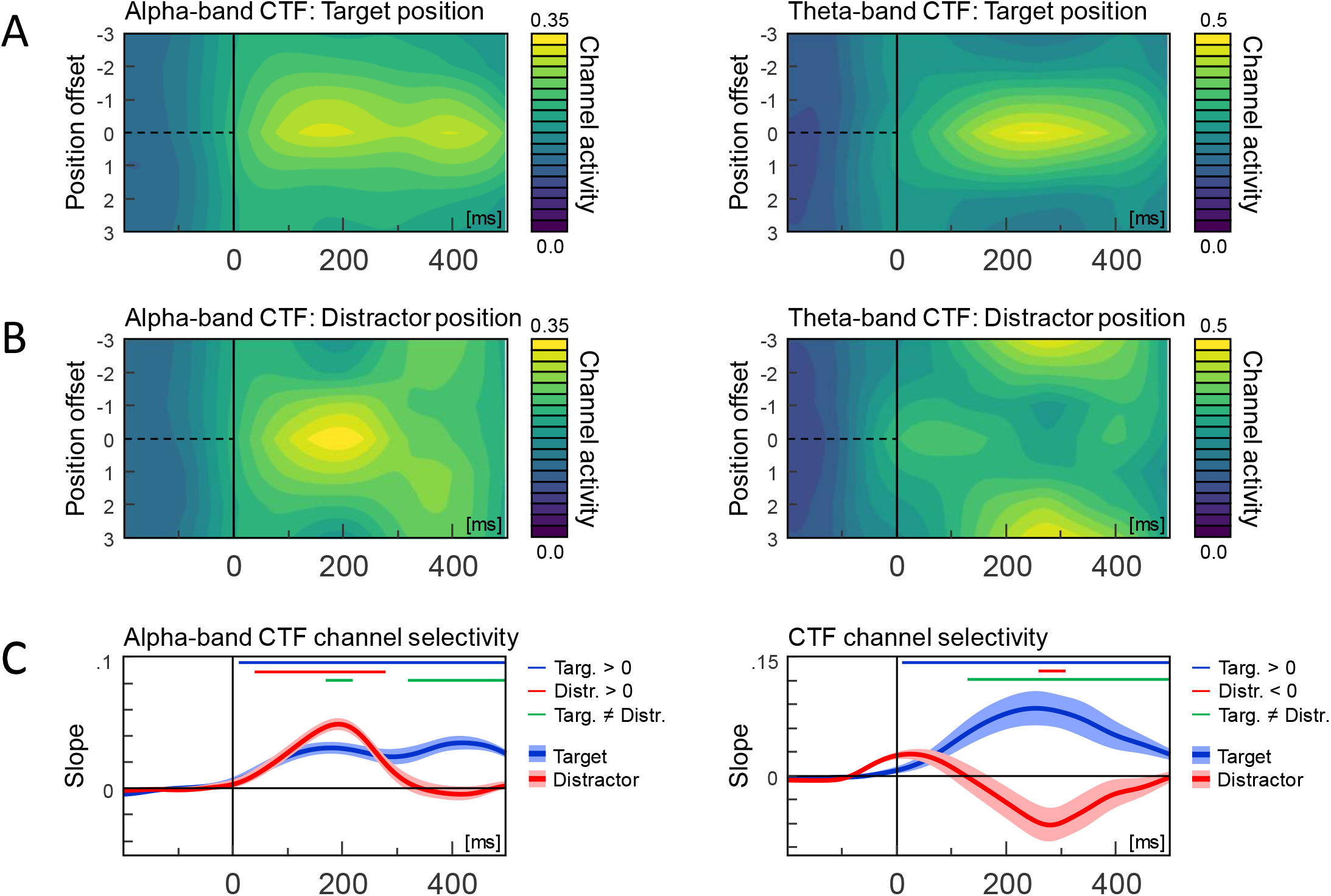
Channel tuning functions for evoked alpha-band (8-12 Hz) activity (left panels) and evoked theta-band (4-8 Hz) activity (right panels). All CTFs are trained on target position in distractor-absent trials and tested on (A) target position and (B) distractor positions in distractor-present trials. The slope of the CTFs in (A) and (B) is show in (C) as blue lines (target) and red lines (distractor). Thin blue lines show time point where the target-CTF slope is different from zero, thin red lines show where the distractor-CTF is different from zero. Thin green lines show where target-CTF and distractor-CTF differ from one another. Only time points in clusters of at least 50 subsequent time points (= 50 ms) with p < .05 are highlighted. Shades indicate standard errors of the mean, corrected for individual differences (Cousineau, 2005).

The IEM run on theta bandpass (4-8 Hz) filtered data shows a slope reliably different from zero from 5-500 ms for the target stimulus; thus the spatial distribution of activity in the theta band tracked the position of the target. Theta band activity also covaried with distractor position from 262-312 ms, but in this case the lowest level of channel activity was at the position of the distractor, such that there was an inverted tuning function surrounding the distractor position. Given that the target and distractor positions were completely balanced in this analysis, the inverted tuning function observed in the distractor analysis suggests a spatially-focused suppression of activity at the distractor position.

## Discussion

The key finding of the present study was to document a spatial gradient of suppression surrounding salient distractors while observers engaged in visual search. We found that with decreasing distance between target and distractor in a visual search task, both the degree of distractor interference and the Pd, a neural marker of active suppression, increased in amplitude. Our results suggest that more suppression is applied when salient, irrelevant information is presented in close proximity to a target stimulus. At the same time, when target and distractor positions coincided and spatially directed suppression could interfere with target processing, the Pd was eliminated.

We used a novel additional singleton paradigm that was different from classical search tasks in that target and distractor location were independently positioned, allowing them to occupy the same location in a subset of trials. We replicated earlier findings showing that a salient distractor results in RT costs (Theeuwes, 1992, 2010; Feldmann-Wüstefeld & Schubö, 2013; Sawaki & Luck, 2013). RT costs were most pronounced for singletons adjacent to the target and monotonically decreased to more distant locations, replicating earlier findings (Caputo & Guerra, 1998; Gaspar & Mcdonald, 2014; Koch et al., 2013; Kwak et al., 1991; Mounts, 2000; Starreveld et al., 2004; Theeuwes et al., 2004). This gradient of RT costs implies a spatial form of interference. Experiment 1 showed RT costs and distance effects when the target was the additional singleton at chance level (1/6 of trials), i.e., when there was no incentive for participants to attend red. Experiments 2 and 3 replicated the results with triple compound stimuli, i.e. when an additional red ring was presented around the target in 1/6 of the trials. Finally, Experiment 4 showed the same pattern of results when the distractor color was determined by whether the distractor shared or did not share its location with the target, showing that salient distractors interfere with target processing regardless of whether they are linked to relevant or irrelevant locations. Interestingly, all four experiments showed that in trials in which target and distractor shared the same location, RTs were shorter than for any other distance, but longer than for distractor-absent trials. This empirical pattern provides further evidence that some of the interference from singleton distractors is not explained by spatial capture alone.

Neural markers of attention provided further insight into how selection and suppression varied across these experimental conditions. We replicated prior work showing that the target-N2pc is smaller when target and distractor are in close proximity (Gaspar & McDonald, 2014), suggesting that attentional selection of relevant information may be impaired by nearby distractors. Using a systematic lateralization approach also allowed us to isolate specific neural signatures of distractor processing (Hickey et al., 2009). Distractors elicited an N2pc, followed by a distractor positivity (Pd), suggesting they captured attention and were then suppressed (Burra & Kerzel, 2013; Feldmann-Wüstefeld & Schubö, 2013; Sawaki & Luck, 2013). The Pd component was larger for distractors adjacent to a target than for distractors further away. Thus, both the Pd as a measure of distractor suppression and the N2pc as a measure of target enhancement were affected by target-distractor distance, but in an inverse fashion. This is in line with the neural ambiguity account that posits increased interference for stimuli that are presented in the same receptive field (Desimone & Duncan, 1995; Hickey & Theeuwes, 2011; Luck et al., 1997). Our results suggest that the visual system resolves neural ambiguity by applying more suppression to distractors near the target at the cost of enhancing the target itself. It was previously suggested that increased RT costs for distractors near targets are due to more attentional capture (Hickey & Theeuwes, 2011; Mounts, 2000). However, our results were inconsistent with this hypothesis, because the N2pc elicited by the distractor did not vary as a function of distance. Although Pd increased with target-distractor proximity, no Pd was observed when they shared the same location, and instead a large N2pc was found. This suggests that target-distractor compounds were not suppressed but rather were strongly attended, in line with the notion that two coinciding singletons produce high activations on the priority map due to a redundancy gain (Krummenacher et al., 2002) and due to top-down and bottom-up factors dovetailing (Fecteau & Munoz, 2006; Itti & Koch, 2001; Wolfe, 2007). The Ppc that was associated with early suppression (Barras & Kerzel, 2017; Tobias Feldmann-Wüstefeld & Vogel, 2019; Weaver et al., 2017) was not affected by target-distractor distance. This may be because the Ppc reflects an early “attend-to-me” signal (Sawaki & Luck, 2010) signal that is not modulated by the amount of interference but rather reflects activity on the priority/salience map (Barras & Kerzel, 2017). In this context the Pd can be seen as a response to that signal that reflects increasing levels of suppression of the distractor as interference from the distractor increases.

The IEM allowed us to examine spatially selective EEG activity that tracked the position of both targets and distractors. Alpha-band (8-12 Hz) activity showed that initially, both targets and distractors were attended, but distractors more so than targets. The color contrast was strong for the distractor singletons, such that they were likely more salient than the shape-defined target singletons. Thus, stronger initial capture by the distractor falls in line with the notion that salient stimuli capture attention automatically and may receive priority over less salient, yet task-relevant items (Theeuwes, 2010; see also Wyble et al., in press). The spatial selectivity for distractor positions was transient, however, and later exceeded by the spatial selectivity of targets (from ~280 ms on, only targets were attended), in line with the idea that attentional capture is followed by rapid disengagement (Belopolsky et al., 2010) as well as with the idea that top-down attention deployment eventually overrides attentional capture and thus supports encoding of task-relevant information (Leber & Egeth, 2006). We also show that theta- and alpha-band activity are picking up on qualitatively different aspects of attentional processing. Although theta-band (4-8 Hz) activity tracked the position of the target, it showed below-baseline spatial selectivity for the distractor, most pronounced at ~290 ms. Channel activity in the theta band showed a graded tuning profile, with the lowest channel activity observed at the position of the distractor, and a monotonic rise in channel activity as distance from that position increased. Thus, spatially selective activity in the theta band reveals a relative suppression of channel activity at distractor positions, indicating a spatially-directed form of distractor suppression.

### Spatial and nonspatial interference

The increasing RT costs with decreasing target-distractor distance indicate that distractors induced spatial interference. The graded Pd and target-N2pc amplitude as well as the graded tuning profile of alpha- and theta-band activity further support this notion. Analogously, the reduced RT costs and the larger target-N2pc for targets and distractors sharing the same location compared to dist>0 trials indicate a spatial *benefit*. We also found compatibility effects: RTs were faster when target and distractor identity were identical, indicative of spatial attention deployment towards the distractor (Becker, 2007; Folk & Remington, 2006). At the same time, distractor-N2pc amplitudes were distance-invariant, demonstrating that distractors did not capture attention more when they were near targets. On the contrary, when distractor and target coincided, the N2pc was larger than for distractors or targets presented alone. Interestingly, performance was worse in this condition than when no distractor was presented at all (distance-zero trials), even though a pure spatial capture account (Theeuwes, 1992, 2010) would predict that spatially coincident targets and distractors should elicit robust spatial capture and thus produce the shortest RTs. Thus, distractor interference for distance-zero trials suggests that distractors also induced non-spatial filtering costs, in line with prior demonstrations (Becker, 2007; Folk & Remington, 1998; Folk & Remington, 2006; Wykowska & Schubö, 2011). Same location costs were previously observed in contingent capture paradigms in which a salient distractor was occasionally presented at the upcoming target position (Carmel & Lamy, 2014; Eimer, Kiss, Press, & Sauter, 2009; Lamy, Leber, & Egeth, 2004; Schoeberl, Ditye, & Ansorge, 2018). One possibility is that target and distractor compete for encoding resources, such that the extraction of target features is delayed even when target and distractor are presented in the same location (Folk & Remington, 2006; Kahneman et al., 1983; Treisman et al., 1983).

We used event-related potentials (ERPs) and an inverted encoding model (IEM) to demonstrate that the visual system responds to distractor interference with a spatial gradient of suppression: The Pd component in the ERP increased with increasing proximity of target and distractor while the theta-band IEM showed most suppressed activity at the distractor location and monotonically decreasing suppression towards more distant locations. Whereas suppression is strongest for distractors presented near the target, selection is weakest for targets presented near a distractor. In sum our results provide robust evidence for a spatially directed process for suppressing distractor interference during goal-driven visual search.

